# Motto: Representing motifs in consensus sequences with minimum information loss

**DOI:** 10.1101/607408

**Authors:** Mengchi Wang, David Wang, Kai Zhang, Vu Ngo, Shicai Fan, Wei Wang

**Author notes:** M. Wang and D. Wang contributed equally to this work.

## Abstract

Sequence analysis frequently requires intuitive understanding and convenient representation of motifs. Typically, motifs are represented as position weight matrices (PWMs) and visualized using sequence logos. However, in many scenarios, representing motifs by wildcard-style consensus sequences is compact and sufficient for interpreting the motif information and search for motif match. Based on mutual information theory and Jenson-Shannon Divergence, we propose a mathematical framework to minimize the information loss in converting PWMs to consensus sequences. We name this representation as sequence Motto and have implemented an efficient algorithm with flexible options for converting motif PWMs into Motto from nucleotides, amino acids, and customized alphabets. Here we show that this representation provides a simple and efficient way to identify the binding sites of 1156 common TFs in the human genome. The effectiveness of the method was benchmarked by comparing sequence matches found by Motto with PWM scanning results found by FIMO. On average, our method achieves 0.81 area under the precision-recall curve, significantly (*p*-value < 0.01) outperforming all existing methods, including maximal positional weight, Douglas and minimal mean square error. We believe this representation provides a distilled summary of a motif, as well as the statistical justification.

**AVAILABILITY:** Motto is freely available at http://wanglab.ucsd.edu/star/motto.

## 1. INTRODUCTION

Motif analysis is crucial for uncovering sequence patterns, such as protein-binding sites on nucleic acids, splicing sites, epigenetic modification markers and structural elements^1^. A motif is typically represented as a Position Weight Matrix (PWM), in which each entry shows the occurrence frequency of a certain type of nucleic acid at each position of the motif. PWMs are often visualized by sequence logo^1^, which requires a graphical interface. Recently, several studies have shown the usefulness of representing motifs using kmers^2–5^; despite the power of this representation in machine learning models, it is cumbersome to have a set of kmers to characterize a single motif. In many scenarios, motifs can be sufficiently represented by regular expressions of the consensus sequences, such as [GC][AT]GATAAG[GAC] for the GATA2 motif. This representation is the most compact and intuitive way to delineate a motif. In the GATA2 motif example, the GATAAG consensus in the center is the most prominent pattern that would be read off the PWM or sequence logo. For this reason, consensus sequences are still widely used by the scientific community. Sequence pattern in the regular expression is used to highlight motif occurrence in popular genome browsers such as UCSC^6^ and IGV^7^. Consensus sequences are assigned to *de novo* motifs and sequences for informative denotations^8–11^. Wildcard-like sequence patterns are also supported in DNA oligo libraries synthesis by major vendors including Invitrogen, Sigma-Aldrich, and Thermo-Fisher.

However, current methods that convert PMWs to consensus sequence are often heuristic. One simple approach is taking the nucleotide with maximal frequency at each position to define the consensus sequence (eg. GGTCAAGGTCAC for ESRRB). Unsurprisingly, this could misrepresent positions with similar frequencies (eg. 0.26, 0.25, 0.25, 0.24, which should have been assigned as N). Alternatively, Douglas *et al.*^*12*^ proposed in 1987 to follow a set of rules: use the single nucleotide with the highest frequency when it exceeds 0.50 and two times the second highest frequency; else, use the top two dinucleotides when their total frequencies exceed 0.75; else, use N. However, these rules are arbitrary, inflexible and lack a mathematical framework.

Here we present Motto, a sequence consensus representation of motifs based on information theory and ensures minimal information loss when converted from a PWM (**Figure 1**). We provide a standardized solution that determines the optimal regular expression of motif consensus sequence. We have also implemented an lightweight and easy-to-use Python package with versatile options for the biologists.

**Figure 1.**
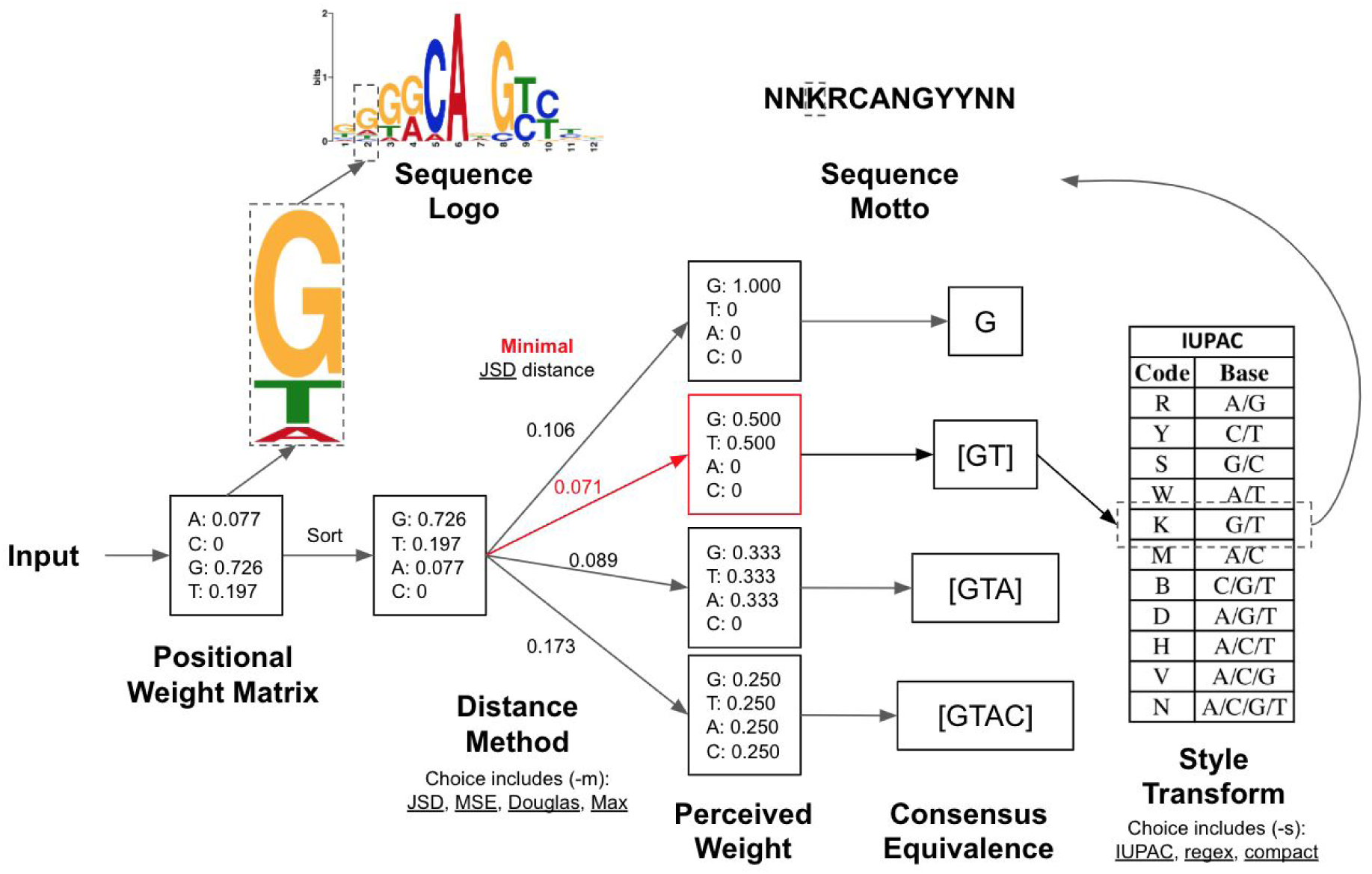
Overview of sequence Motto and comparison with sequence logo. Given a motif PWM as the input, Motto outputs a consensus that minimizes information loss. Here we show how the sequence Motto of the human transcription factor P73 is determined through the minimal JSD method.

## 2. METHODS

### Problem formulation

A positional weight matrix (PWM) maps *I* × *P* → ℝ, where *I* is the set of indices of motif positions, and *P* is the frequency of the nucleotide in the motif. For a given position *i* ∈ *I*, let *C*(*i*) denote the perceived frequencies for a combination of nucleotides, defined by equal frequencies shared among included nucleotides. For example, a *C*(*i*) of [ACT] has the frequencies of [0.333, 0.333, 0, 0.333] for [A, C, G, T], respectively. Thus, we consider the optimal consensus sequence as a series of combination of nucleotides that has the most similarity between *C*(*i*) and *P* (*i*) for each position *i* ∈ *I*.

### Minimal Jensen-Shannon Divergence (JSD) method

Here, we propose to use Jensen-Shannon Divergence (JSD) to measure the similarity between *C*(*i*) and *P* (*i*). JSD has been widely used in information theory to characterize the difference between distributions^13^. Using this metric, the combination of nucleotides with the least JSD from *C*(*i*) to *P* (*i*) will have the minimal “information loss”, and is thus considered as the optimal consensus nucleotide.

To efficiently compare JSD between all possible nucleotide combinations, we propose the following algorithm (**Figure 1**). Given a motif in its PWM form, having *k* positions, and *n* possible elements at each position, then the probability of *j* th element at *i* th position is given by *P* (*i, j*), where 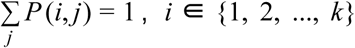, and *j* ∈ {1, 2, …, *n*}. First, we sort the elements of the PWM in descending order, so that:

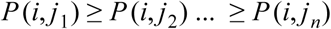

For example, at the 2nd (*i* = 2) position of the human transcription factor P73 (**Figure 1**), the nucleotides are sorted by occurrence frequencies so that:

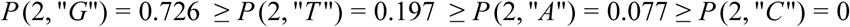

Next, we denote *m* as the number of different elements to be presented in the output consensus at the *i* th position, *m* ∈ {1, 2, …, *n*}. If a nucleotide is contained in a position in the consensus sequence, all the other nucleotides with higher frequencies at the *i* th position must also be included. Therefore, identification of the optimal consensus sequence is equivalent to identifying the optimal *m*.

When represented by the consensus sequence Motto, each nucleotide is considered as equally probable at a given position. For a position with {*j*_*i*,1_, *j*_*i*,2_, …, *j*_*i,m*_} nucleotides, *C*(*i*, 1) = *C*(*i*, 2) = … = *C*(*i, m*) = 1/*m*. The closer this distribution is to the original distribution of nucleotide frequencies, the better approximation of the consensus motif is to the original PWM. The optimal *m* (denoted as *m*\*) can be determined by minimizing the JSD between the two distributions:

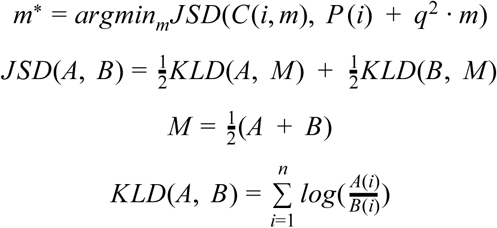

Here, *q* ∈ [0, 1] is the ambiguity penalty, a parameter input from the user to penalize large value of *m* in the output. When *q*=0, the optimal *m*\* marks the canonical minimal JSD. When *q*=1, *m*\* is guaranteed to be 1, hence the output consensus nucleotide is {*j*_1_} (equivalent to using nucleotide with the maximal frequency). Thus, the optimal consensus nucleotide at the *i*th position is:

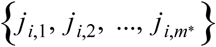

Repeat this procedure for every position *i* ∈ {1, 2, …, *k*}, the final optimal consensus sequence is given by:

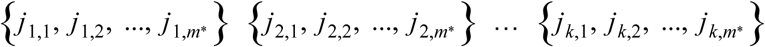

### Minimal Mean Squared Error (MSE) method

For comparison purposes, we have also implemented minimal mean squared error (MSE) method, which is another widely-used metric to measure distribution discrepancy^14^. The rest of the implementation is unchanged, except for the optimal *m* (*m*\*) is now determined by minimizing the MSE between the two distributions:

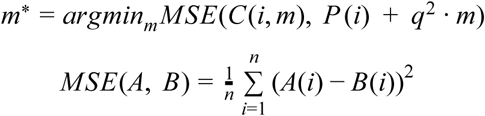

### Evaluating motif occurrence sites

We have collected 1156 common transcription factors from human and mouse from the databases of Transfac^15^, Jaspar^16^, Uniprobe^17^, hPDI^18^, and HOCOMOCO^19^. Each PWM is converted into consensus sequences, using default options of the four discussed methods: JSD (described above), MSE (described above), Douglas^12^ and the naive approach of using the maximal frequency. Motif occurrence sites are determined in the human genome (hg19), matched by their regular expressions. The ground truth of the occurrence sites is determined by scanning the original PWMs with FIMO^20^ using a 1e-5 *p*-value cutoff. The resulting *p*-values are converted into a significance score (-log(*p*-value)) and assigned to the matched motif occurrence sites from sequence Mottos. Thus, the area under the precision-recall curves^21^(auPRC) is calculated by comparing the motif occurrence sites and their significance scores. Resulting auPRCs are averaged and a paired (by each motif) t-test is conducted to determine performance. Comparisons with significance (*p*-value < 0.01) are shown (**Figure 3**).

**Figure 2.**
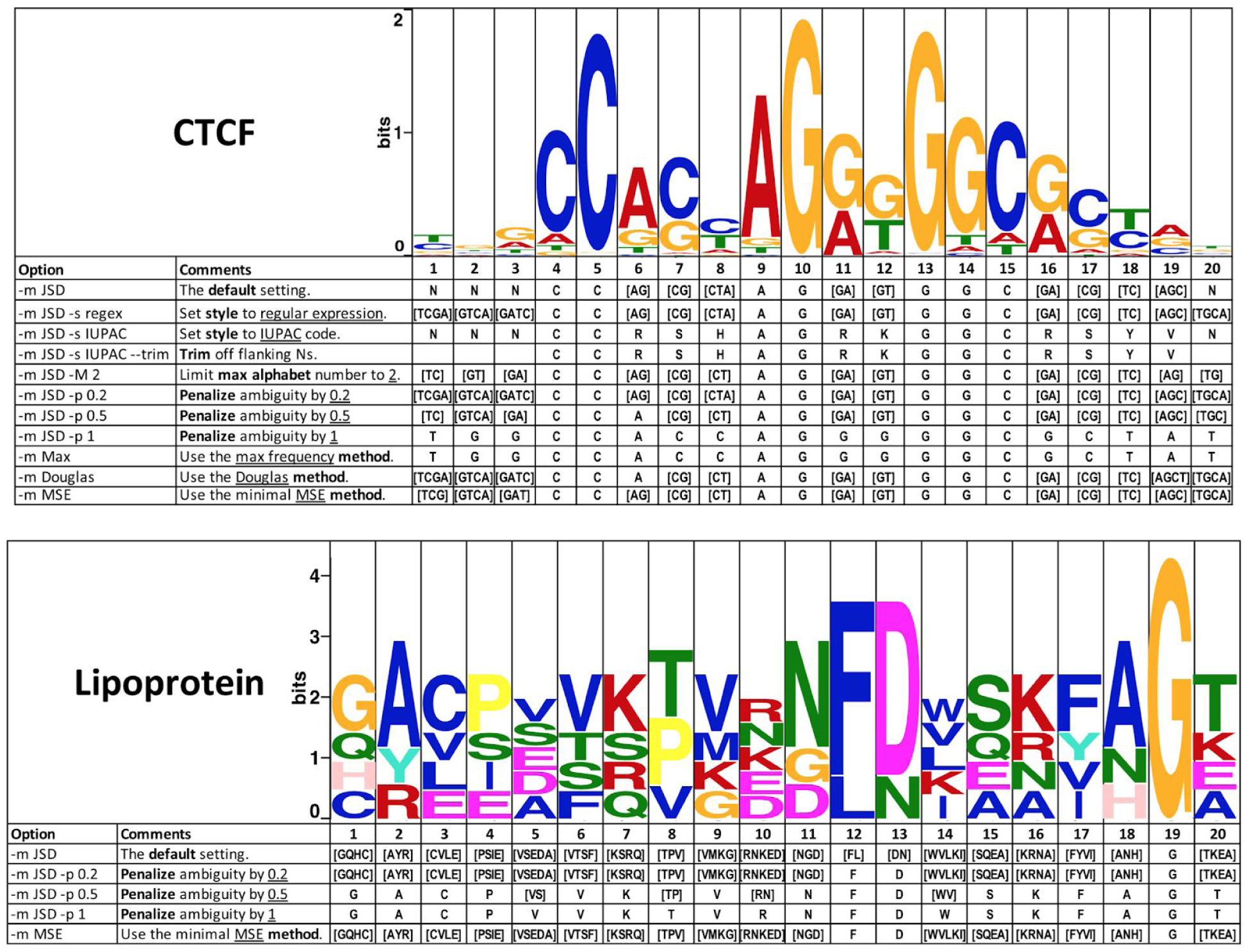
Example usage using human CTCF (**upper panel**) and lipoprotein binding sites from Bailey 1994^24^ (**lower panel**). The original PWM is shown in sequence logo. Different Motto options resulted in various consensus sequence output at each position. In particular, “-m/--method” specifies the method: JSD (default), MSE (minimal mean square error), Douglas^12^, or Max (using maximal frequency at each position); “-s/--style” specifies the output style: IUPAC^23^ (single alphabet for nucleotide combinations), regex (regular expression), or compact (convert [ACGT] to N in regex); “-t/--trim” is an option for trimming off the flanking Ns; “-p/--penalty” specifies a weight between 0 to 1 that penalizes ambiguity at each position. (For details see **Methods**)

**Figure 3.**
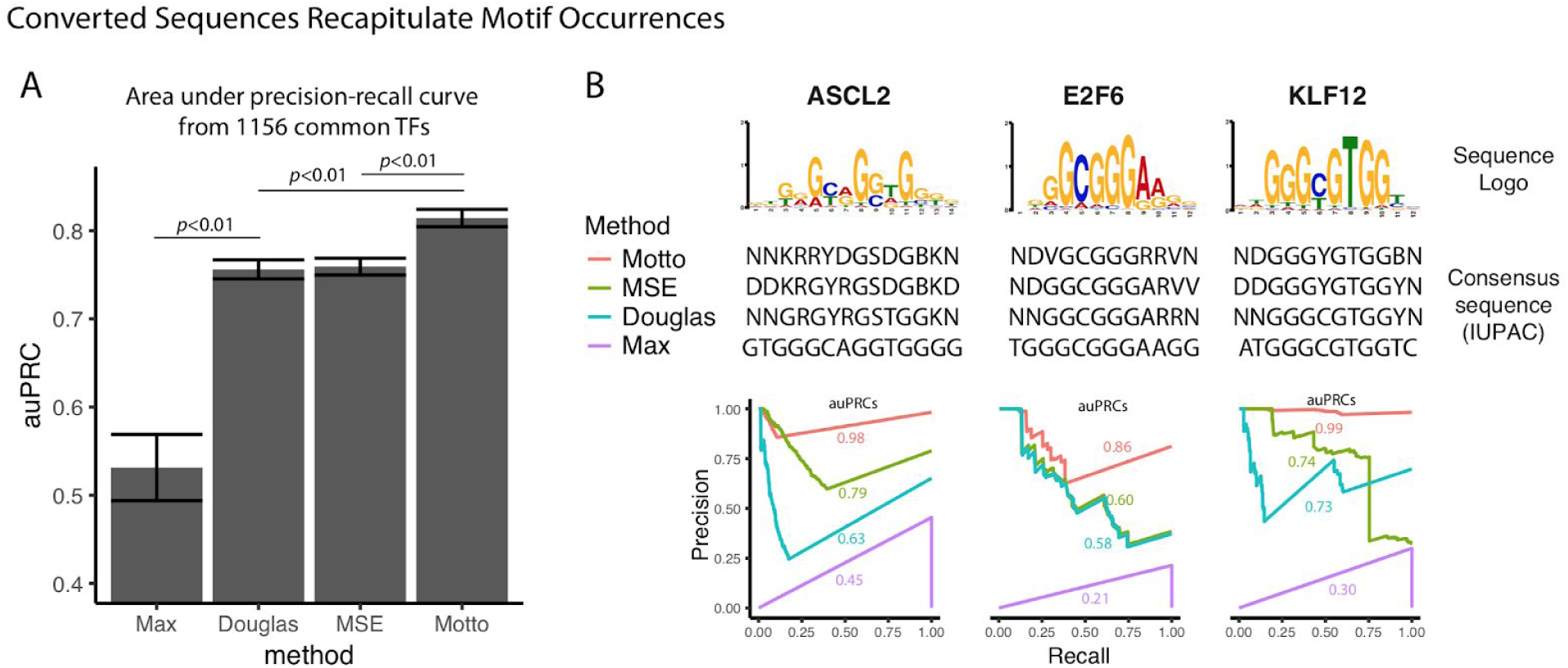
Converted sequence Mottos recapitulate motif occurrence sites of 1156 common human and mouse transcription factors (TFs) in the human genome (hg19). **A**. Averaged area under the precision-recall curve (auPRC) using Motto (default method with minimal JSD) compared with existing alternative methods. *P*-value determined by paired t-test. **B**. Comparison in three example TFs showing the differences of consensus sequences (shown in IUPAC^23^ coding for better alignment) and performances.

## 3. FEATURES AND EXAMPLES

Motto takes the MEME format of PWM as the input because of its popularity. The MEME format is supported by the majority of the motif databases^19^, and the MEME suite provides packages for integrative analysis and conversion from other motifs formats^10^. The recently proposed kmer-based motif models also support conversion to MEME format^2,3^. Our package is lightweight and open-source. The algorithm is efficiently implemented in python and the conversion for 1000 motif sequences typically takes less than two seconds. In addition, perhaps expectedly, downstream analysis like matching motif occurrences using the regular expression of sequence Motto is much faster (about 5 seconds for a common PWM on a chromosome, implemented in-house with python) than a conventional PWM scanning (about 1 minute, scanned with FIMO^20^). By default, the Motto package takes a motif in the MEME format, parses the header to get the nucleotide, computes the optimal consensus sequences based on the minimal JSD method, and then outputs the sequence in a compact format (**Figure 1**). Motto provides flexibility at each step along this process. Input can be from a file, or from standard input, and Motto can consider nucleotides, amino acids and customized alphabets such as CpG and non-CpG methylation^22^. Four methods are made possible for comparison: maximal probability (Max), heuristic Douglas method (Douglas), minimal mean squared root (MSE) method, and our proposed minimal Jensen-Shannon Divergence (JSD) (See **Methods**). Three output styles are provided: (1) IUPAC uses a single alphabet to represent the combination of nucleotides (eg, S for [CG]) and is the most compact form, but require reference to the nomenclature^23^; (2) regular expression (“regex”) enumerate all output consensus nucleotide ranked by occurrences and is recommended for downstream analysis, such as motif occurrence and oligo designs; (3) “compact” (the default) is the same as “regex”, except that it replaces [ACGT] with N. To trim off Ns ([ACGT]s) at both ends of the output sequences, a optional flag “--trim” is provided. If the users prefer consensus with more certainty (eg, prefer [AC] to [ACG]), they can use either “--maxAlphabet” as a hard limit to the number of alphabets allowed, or use “--penalty” to penalize ambiguity (see **Methods**).

Effects of these options are shown using an example of human transcription factor CTCF (**Figure 2**, **upper panel**). Unsurprisingly, MSE, JSD, and Douglas are more representative than the naive maximal probability methods. For example, position 1,2,3 and 20 with low information content (<0.2) in CTCF, are justifiably called as “N” by JSD and Douglas, which is an improvement over strictly calling the top nucleotide. MSE considers [TCG] and [GAT] more representative at the 1st and the 3rd position but agrees with JSD and Douglas at the 2nd and 20th. Similarly, JSD, MSE, and Douglas successfully capture strong double-consensus patterns at indices 7, 11, 12 and 16, which maximal probability fails to capture. The advantage of JSD over Douglas is noticeable at index 6, where the logo of CTCF shows a dominating AG consensus. While JSD finds this co-consensus, Douglas disregards G that barely misses the cutoff. In addition, at index 19, the logo of CTCF shows a strong three-way split among A, G and C, but Douglas, by its rules (as described previously), ignores all such triple patterns. In addition, among the four methods, only the JSD and MSE are capable of generating consensus sequences for amino acid motifs^24^ (**Figure 2**, **lower panel**). Due to its arbitrary nature, heuristic methods like Douglas have difficulties defining decision boundary for motifs more than 4 nucleotides. In such cases, JSD and MSE provide more mathematically rigorous information than Douglas and oversimplified maximal consensus methods. With increased penalty level at 0, 0.2, 0.5 and 1 respectively, the consensus sequence smoothly progresses towards single nucleotide consensus (**Figure 2**). Such flexibility gives an advantage to users that are biased towards more defined consensus results.

To quantify how well these four methods summarize the information in the original PWMs, we converted 1156 common human and mouse transcription factors to consensus sequences and compared their matched occurrences (by regular expression) in the human genome (hg19) with conventional motif sites scanned by FIMO^20^ with PWMs, which is how conventionally motif sites are determined (see **Methods**). We observe that using the JSD method has resulted in the best (0.81±0.01) area under the Precision-Recall curve (auPRC), significantly (*p*-value < 0.01) when compared with existing alternative methods, including MSE (0.76±0.01), Douglas (0.76±0.01), and maximal frequency (0.53±0.04) (**Figure 2**).

In summary, Motto provides a mathematical framework and a set of convenient features to epitomize PWMs in a compact, intuitive and informative manner.

## AUTHOR CONTRIBUTIONS

Mengchi Wang conceived the idea, implemented the package, performed the analyses, and wrote the manuscript; David Wang implemented key aspects of the package, contributed to the data analysis and manuscript preparation; Kai Zhang, Vu Ngo, and Shicai Fan contributed to data analysis and manuscript preparation; Wei Wang supervised the analyses and the project.

## REFERENCE

1. Schneider, T. D. & Stephens, R. M. Sequence logos: a new way to display consensus sequences. Nucleic Acids Res. 18, 6097–6100 (1990).

2. Fletez-Brant, C., Lee, D., McCallion, A. S. & Beer, M. A. kmer-SVM: a web server for identifying predictive regulatory sequence features in genomic data sets. Nucleic Acids Res. 41, W544–56(2013).

3. Ghandi, M., Lee, D., Mohammad-Noori, M. & Beer, M. A. Enhanced regulatory sequence prediction using gapped k-mer features. PLoS Comput. Biol. 10, e1003711 (2014).

4. Guo, Y., Tian, K., Zeng, H., Guo, X. & Gifford, D. K. A novel k-mer set memory (KSM) motif representation improves regulatory variant prediction. Genome Res. 28, 891–900 (2018).

5. Zeng, H., Hashimoto, T., Kang, D. D. & Gifford, D. K. GERV: a statistical method for generative evaluation of regulatory variants for transcription factor binding. Bioinformatics 32, 490–496 (2016).

6. Kent, W. J. et al. The human genome browser at UCSC. Genome Res. 12, 996–1006 (2002).

7. Robinson, J. T. et al. Integrative genomics viewer. Nat. Biotechnol. 29, 24 (2011).

8. Wang, M. et al. Identification of DNA motifs that regulate DNA methylation. bioRxiv 573352 (2019). doi:10.1101/573352

9. Whitaker, J. W., Chen, Z. & Wang, W. Predicting the human epigenome from DNA motifs. Nat. Methods 12, 265–72, 7 p following 272 (2015).

10. Bailey, T. L. et al. MEME SUITE: tools for motif discovery and searching. Nucleic Acids Res. 37, W202–8 (2009).

11. Heinz, S. et al. Simple combinations of lineage-determining transcription factors prime cis-regulatory elements required for macrophage and B cell identities. Mol. Cell 38, 576–589 (2010).

12. Cavener, D. R. Comparison of the consensus sequence flanking translational start sites in Drosophila and vertebrates. Nucleic Acids Res. 15, 1353–1361 (1987).

13. Lin, J. Divergence measures based on the Shannon entropy. IEEE Trans. Inf. Theory 37, 145–151 (1991).

14. Lele, S. Euclidean Distance Matrix Analysis (EDMA): Estimation of mean form and mean form difference. Math. Geol. 25, 573–602 (1993).

15. Matys, V. et al. TRANSFAC® and its module TRANSCompel®: transcriptional gene regulation in eukaryotes. Nucleic Acids Res. 34, D108–D110 (2006).

16. Portales-Casamar, E. et al. JASPAR 2010: the greatly expanded open-access database of transcription factor binding profiles. Nucleic Acids Res. 38, D105–10 (2010).

17. Robasky, K. & Bulyk, M. L. UniPROBE, update 2011: expanded content and search tools in the online database of protein-binding microarray data on protein–DNA interactions. Nucleic Acids Res. 39, D124–D128 (2011).

18. Xie, Z., Hu, S., Blackshaw, S., Zhu, H. & Qian, J. hPDI: a database of experimental human protein–DNA interactions. Bioinformatics 26, 287–289 (2010).

19. Kulakovskiy, I. V. et al. HOCOMOCO: towards a complete collection of transcription factor binding models for human and mouse via large-scale ChIP-Seq analysis. Nucleic Acids Res. 46, D252–D259 (2018).

20. Grant, C. E., Bailey, T. L. & Noble, W. S. FIMO: scanning for occurrences of a given motif. Bioinformatics 27, 1017–1018 (2011).

21. Davis, J. & Goadrich, M. The Relationship Between Precision-Recall and ROC Curves. in Proceedings of the 23rd International Conference on Machine Learning 233–240 (ACM, 2006).

22. Ngo, V., Wang, M. & Wang, W. Finding de novo methylated DNA motifs. Bioinformatics (2019). doi:10.1093/bioinformatics/btz079

23. Johnson, A. D. An extended IUPAC nomenclature code for polymorphic nucleic acids. Bioinformatics 26, 1386–1389 (2010).

24. Bailey, T. L., Elkan, C. & Others. Fitting a mixture model by expectation maximization to discover motifs in bipolymers. (1994).

